# AnchoRNA: Full virus genome alignments through conserved anchor regions

**DOI:** 10.1101/2025.01.30.635689

**Authors:** Tom Eulenfeld, Sandra Triebel, Peter F. Stadler, Manja Marz

**Affiliations:** RNA Bioinformatics and High-Throughput Analysis, Friedrich Schiller University Jena, Germany; European Virus Bioinformatics Center, Jena, Germany; Cluster of Excellence Balance of the Microverse, Friedrich Schiller University Jena, Germany; Bioinformatics Group, Department of Computer Science & Interdisciplinary Center for Bioinformatics, Leipzig University, Germany; Max Planck Institute for Mathematics in the Sciences, Leipzig, Germany; German Centre for Integrative Biodiversity Research (iDiv) Halle-Jena-Leipzig, Germany; Department of Theoretical Chemistry, University of Vienna,Austria; Facultad de Ciencias, Universidad National de Colombia, Bogotá, Colombia; Center for Non-coding RNA in Technology and Health, University of Copenhagen, Denmark; Santa Fe Institute, USA; Fritz Lipmann Institute-Leibniz Institute on Aging, Jena, Germany

**Keywords:** Multiple sequence alignment, Conserved regions, Viral genomes, Anchors

## Abstract

**Background:** Multiple sequence alignment of full viral genomes can be challenging due to factors such as long sequences, large insertions/deletions (spanning several 100 nucleotides), large number of sequences, sequence divergence, and high computational complexity in particular when computing alignments based on RNA secondary structures. Standard alignment methods often face these issues, in particular when processing highly variable sequences or when specific phylogenetic analysis is required on selected subsequences.

**Results:** We present an algorithm to determine high quality anchors that define partitions of sequences and guide the alignment of viral genomes to respect well conserved, and therefore functionally significant, regions. This new approach is implemented in the Python-based command line tool `AnchoRNA`, which is designed to identify conserved regions, or anchors, within coding sequences. By default, anchors are searched in translated coding sequences accounting for high mutation rates in viral genomes. `AnchoRNA` enhances the accuracy and efficiency of full-genome alignment by focusing on these crucial conserved regions. AnchoRNA anchors are compared with those produced by three other anchor-finding tools, and anchor-guided alignments are evaluated using three alignment programs. Utilizing a dataset of 55 representative *Pestivirus* genomes, `AnchoRNA` identified 55 anchors that are used for guiding the alignment process. The incorporation of these anchors led to improvements across tested alignment tools, highlighting the effectiveness of `AnchoRNA` in enhancing alignment quality, especially in viral genomes.

**Conclusions:** We demonstrate that `AnchoRNA` effectively improves the alignment of complete viral genomes by identifying conserved anchor regions that guide multiple sequence alignments.

## 1 Introduction

Multiple sequence alignment (MSA) is a crucial step in biological sequence analysis, serving as the basis for many downstream applications, including functional annotation, evolutionary studies, and structural predictions (e.g. Lamkiewicz et al., 2023). Despite its importance, MSA remains a challenging problem due to its inherent complexity and the diversity of biological sequences, particularly in the context of viral genomes (e.g. Hufsky et al., 2023, 2020; Madhugiri et al., 2014; Hölzer and Marz, 2020). The traditional approach, which focuses on local alignments, has evolved significantly, leading to advancements in alignment accuracy with tools like `ClustalW` (Thompson et al., 1994), `T-Coffee` (Notredame et al., 2000), `MUSCLE` (Edgar, 2004), `MAFFT` (Katoh, 2002), and `DIALIGN` (Morgenstern et al., 1998). These methods have improved the alignment process by integrating local alignment information into global alignment frameworks and incorporating homology data from databases. However, aligning highly variable sequences, such as full viral genomes, still poses significant challenges that require more specialized approaches. Pais et al. (2014) and Zhang et al. (2022) review existing MSA tools and compare their performance using the BAliBASE benchmark (Thompson et al., 1999). To address the challenge of aligning large numbers of sequences, scalable methods such as PASTA (Mirarab et al., 2014), MAGUS (Smirnov and Warnow, 2020), and TWILIGHT (Tseng et al., 2025) employ divide-and-conquer strategies. Despite these advancements, current MSA tools face limitations when handling certain complexities. Specifically, they often struggle with long sequences, large insertions or deletions (as known for e.g. pestiviruses), and highly divergent sequences, which can lead to suboptimal full virus genome alignments or excessive computational demands, e.g., when simultaneously annotating conserved RNA secondary structures with `LocARNA` (Will et al., 2012).

Anchors help by focusing on conserved regions and by improving alignment accuracy. The precise meaning of anchors is not unequivocally defined in the literature. `DIALIGN` is designed to accommodate constraints on pairwise alignments (Morgenstern et al., 2006). The concept of anchors used here originated in synteny analysis and refers to orthologous sequence elements that appear in all input sequences and therefore subdivide multiple sequence alignments into orthologous blocks delimited by the anchors. Velandia-Huerto et al. (2016) and Berkemer et al. (2017) used such anchors to systematically investigate the turnover of repetitive elements and multi-copy genes such as tRNAs. Anchors in these sense are highly conserved local alignment blocks. In genome-wide synteny analysis, 1-1 orthologous protein coding genes are commonly used as anchors. An annotation-free method to detect them on large scales in closely related genomes was introduced by Käther et al. (2025). By partitioning the global alignment problem into independent regions between consecutive anchors, anchors can drastically reduce the computational demands for long sequences. The smaller search space of the sub-problems not only saves computational resources, but also increases the accuracy. Anchors therefore improve alignments in the presence of large insertions and deletions, or high sequence divergence.

Tools with anchors have been developed for (i) very large genomes: `MUMmer` (Delcher et al., 1999), `Mugsy` (Angiuoli and Salzberg, 2010); (ii) handling duplications and genome rearrangements: `Mauve` (Darling et al., 2004), `progressiveMauve` (Darling et al., 2010), `TBA` (Blanchette et al., 2004); (iii) comparative genomics especially in the context of phylogenetics: `LAGAN`, `Multi-LAGAN` (Brudno et al., 2003b), `AVID` (Bray et al., 2002); (iv) precise alignments based on anchor points: `CHAOS/DIALIGN` (Brudno et al., 2003a), `VISTA genome pipeline` (Dubchak et al., 2009); (v) advanced algorithmic techniques, such as graph-based methods or spaced seeds: `ACANA` (Huang et al., 2005), `FSWM` (Leimeister et al., 2018), `OWEN` (Ogurtsov et al., 2002); (vi) extended general multiple sequence alignment: `ClustalW` (Thompson et al., 1994), `T-Coffee` (Notredame et al., 2000), `MAFFT` (Katoh, 2002); and (vii) larger insertions and deletions: `DIALIGN` (Morgenstern et al., 1998; Morgenstern, 1999). Viral genomes are small, compact, and generally less structurally complex compared to the genomes of larger organisms. Tools designed for large genomes which emphasize complex rearrangements and extensive duplications are thus not well-suited for viral genomes. Multiple sequence alignments of viral sequences are commonly computed using tools such as `MAFFT`. However, the accuracy of full genome alignments depends on factors including sequence divergence and the presence of large insertions and deletions. Additional challenges arise when RNA secondary structure is relevant to the analysis (e.g. Madhugiri et al., 2014). For RNA virus genomes, with a length of up to 40000 nt, the focus is on the development of tools that can efficiently handle high mutation rates and accommodate large numbers (typically 50 to 2000) of genomes in a multiple sequence alignment.

Here, we introduce a method for computing accurate anchors, specifically designed for viral genomes within the same taxonomic group, up to the genus level. The method assumes sufficient sequence homology and a conserved genomic organization to identify anchors in a consistent order. `AnchoRNA` produces anchors that are gap-less local alignments. This subdivides the alignment problem, leading to a better performance both in terms of resource consumption and accuracy of the alignment. By focusing on these key conserved regions, `AnchoRNA` addresses the challenges of long viral genomes, large insertions/deletions, and highly divergent sequences, offering a more accurate and efficient solution for aligning both nucleotide and protein sequences.

## 2 Algorithm and Implementation

The `AnchoRNA` algorithm is summarized in Fig. 1, with pseudocode provided in Algorithm 1 in the appendix. The algorithm assumes that there are no genome rearrangements and proceeds in three stages: first, a window-based similarity search is used on the translated coding regions to find short matches as initial candidates. These are then processed by merging overlapping candidates to larger more significant gap-less alignments. In the final phase, inconsistencies between candidates are resolved by removing less significant candidates, resulting in a co-linear sequence of anchors. Denote the input sequences by *S*^(1)^, *S*^(2)^, … *S*^(*n*)^. An *anchor* consists of a set of *n* intervals with the same length, called *flukes*, one in each sequence, see Fig. 1. Each anchor corresponds to a gap-less alignment of the subsequences of *S*^(1)^, *S*^(2)^, … *S*^(*n*)^ defined by the flukes. `AnchoRNA` uses one of the input sequences, say *S*^(1)^, as a reference. We will refer to *S*^(1)^ as the *guiding sequence*. A sliding window with user-defined fixed length *w* on the guiding sequence specifies a candidate fluke 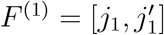. For each sequence *S*^(*k*)^, 2 ≤ *k* ≤ *n*, `AnchoRNA` searches for a matching fluke in a search window [*p*_*k*_, *q*_*k*_] where *p*_*k*_ := *j*_1_ ™ *L* + min(0, *n*_*k*_ ™ *n*_1_) and 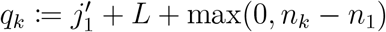 and *n*_*i*_ := |*S*^(*i*)^| denotes the length of sequence *S*^(*i*)^, and *L* is the user-defined `search_range` parameter. That is, if *S*^(1)^ and *S*^(*k*)^ have the same length, the search for a matching fluke is limited to 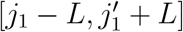. If *S*^(*k*)^ is shorter than the guiding sequence *S*^(1)^, then the search interval is extended by the length difference |*n*_*k*_ ™ *n*_1_| to the left, if it is longer, the interval is extended to the right.

**Figure 1:**
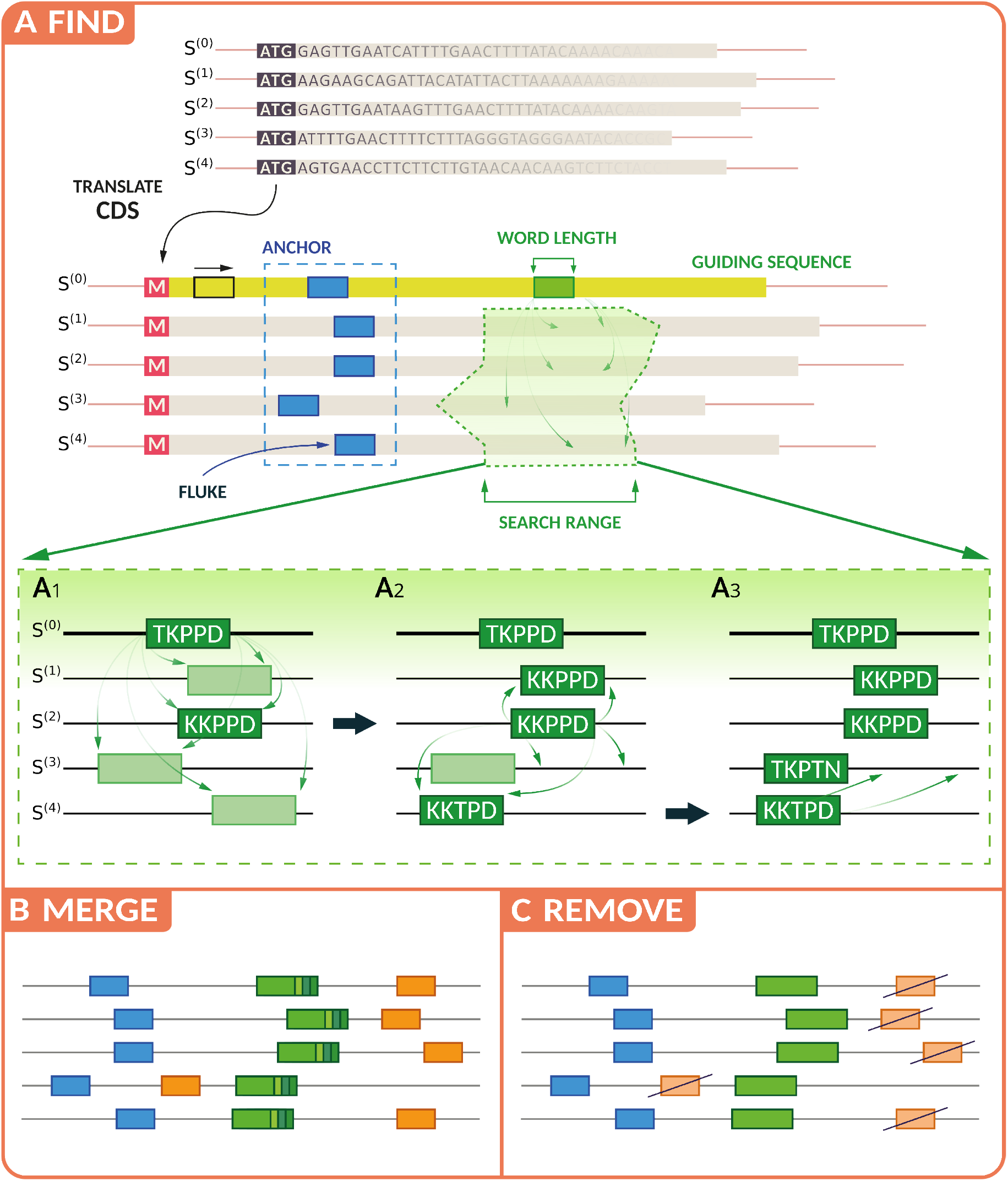
Illustration of the anchor detection algorithm. In this example the translation of the viral RNA sequence is used. Anchor construction is subdivided into three phases: **A** Detection of preliminary anchors starts from a query word in the guiding sequence. For each input sequence, a search window with user-defined width is defined. In the first pass, at most one exact match of the query word in all other sequences in detected. Then, the most similar matches in the remaining sequences are iteratively used as queries. A preliminary anchor consists of one fluke in each sequence, each of which is sufficiently similar to one of the query words. **B** Preliminary anchors are merged if they overlap and their flukes are separated by the same distance in all sequences. The merged anchor candidates are assigned a similarity score. **C** A simple greedy algorithm is used to remove all candidate anchors that still overlap or cross another anchor with higher similarity score. The result is a co-linear sequence of anchors, each of which corresponds to a gap-less local alignment.

`AnchoRNA` attempts to determine an anchor candidate for each position of the guiding sequence, starting with query word *F*_1_ corresponding to the fluke *F* ^(1)^ in *S*^(1)^. The fluke *F* ^(1)^ becomes part of the candidate anchor *A*. The search proceeds then iteratively using an expanding list *Q* of query words starting from *Q* = {*F*_1_}. In each step, the search covers the intervals [*p*_*j*_, *q*_*j*_] of all sequences *S*^(*j*)^ that have not been assigned a fluke in a previous step and determines the word *W* in these intervals with the maximal similarity score *σ*(*W* ) := max_*F*∈*Q*_ *σ*(*W, F* ) over all query words. If *σ* equals or exceeds a threshold *σ*_add_anchor_, then the position of word *W* is added as fluke *F* ^(*j*)^ to the anchor candidate *A*. If *σ* equals or exceeds a threshold *σ*_add_word_ ≥ *σ*_add_anchor_, then *W* is added to the set of query words. By default, the two thresholds are the same. In the first step, therefore, flukes that correspond to identical matches of *F*_1_ are added to *A*, and in later stages, increasingly different words *W* are obtained. The search ends if either all sequences have been assigned a fluke or if the highest-scoring match *W* does not reach the threshold *σ*_add_anchor_, and thus no further fluke can be added to the anchor candidate. The candidate is discarded if the fraction of sequences with flukes does not reach a user-defined threshold. By default only complete anchors, i.e., those with a fluke in every sequence are retained as preliminary anchors.

Each preliminary anchor *A* is associated, by construction, with the set *Q*_*A*_ of query words that were used for its construction. Note that, in general, *Q*_*A*_ contains only a subset of the words *F*_*k*_ defined by flukes *F* ^(*k*)^. The initial query word *W* corresponding to the guiding fluke *F* ^(1)^, however, is always contained in *Q*_*A*_. In order to score the preliminary anchor *A*, we proceed as follows: For each fluke *F* ^(*k*)^ of *A*, we first determine the median 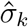 of the similarity scores *σ*(*F* ^(*k*)^, *W* ) with all query words *W* ∈ *Q*_*A*_ and then compute the *anchor score* as the minimum of these median scores over all flukes in the candidate anchor: 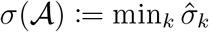.

These initial anchors are processed in two steps. First, anchors *A* and *A*^′^ with overlapping flukes *F* ^(*k*)^ and 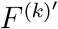 are merged if they are compatible, i.e., the difference between the fluke positions 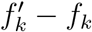 is the same for all *k*. After merging, each anchor consists of flukes 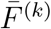 in S^(*k*)^with a common length. The fluke and anchor scores are recomputed after merging.

In the second step anchors are removed to ensure consistency. Note that any overlapping anchors that have not been merged in the previous step necessarily contain contradictory position-wise matches in at least one pair of sequences. Two anchors *A* and *A*^′^ *cross* if the start position of their respective guiding flukes satisfy 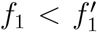 and there is a sequence *S*^(*j*)^ such that the corresponding fluke positions satisfy 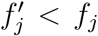. Two preliminary anchors *A* and *A*^′^ are *inconsistent* if they are still overlapping or crossing. A greedy algorithm is then applied to remove *inconsistent* anchors: Starting from the highest scoring preliminary anchor *A*, all preliminary anchors *A*^′^ that are inconsistent with *A* are removed. This step leaves a set of non-overlapping, co-linear anchors, each of which corresponds to a gap-less local alignment. The greedy removal procedure can be restricted to crossing anchors only by setting `aggressive_remove` to false.

Since `AnchoRNA` is designed to address in particular the high levels of sequence divergence commonly observed among viral genomes, the coding RNA sequence is translated to the corresponding amino acid sequence. This feature can also be omitted, e.g. to handle UTR sequences. For amino acid sequences, the `BLOSUM62` (Henikoff and Henikoff, 1992) matrix is used to evaluate the similarity score *σ*, since it accounts for synonymous mutations and mutations for which the corresponding codon is translated to an amino acid with similar properties. Alternatively, the user can choose one of the substitution matrices distributed within the `sugar` package (Eulenfeld, 2025b), which is particularly pertinent when analyzing nucleic acid sequences.

### Complexity and scalability

The time complexity of `AnchoRNA` is dominated by the initial search for preliminary anchors, Figure 1A. The effort depends linearly on the length *n* of the guiding sequence since `AnchoRNA` considers each position as a potential initial fluke for an anchor. Calculating the similarity scores of all words in the search range of the remaining sequences has a complexity of O(*wSN* ), where *S* is the `search_range` parameter *L* plus the maximum of the absolute length differences between the guiding sequence and all other sequences, *w* is the word length of the query, and *N* is the number of input sequences. Finally, updating the best-scoring fluke candidates by a repeated search with new query words results in a runtime that depends quadratically on *N* resulting in worst case complexity of *O*(*nwSN* ^2^). For large *N* ≫ 100, it is advisable to select a representative subset of sequence, see below.

Since `AnchoRNA` needs to store all sequences, the memory consumption amounts to O(*nN* ). It is worth noting that the searches for matching flukes at different position *j*_1_ of the guiding sequence are independent of each other and thus allow a straight-forward parallelization using multiple CPU cores (command line flag `--njobs`) leading to a linear reduction in runtime as the number of cores increases.

### Implementation and Usage

`AnchoRNA` is implemented as a command line tool in python and makes use of the `sugar` (Eulenfeld, 2025b) package, a lightweight toolkit for manipulating sequence and annotation data. The behavior of `AnchoRNA` is affected by several parameters, most importantly the word length *w*, the size *L* of the flanking region defining the search range, the score cutoff for word matches, and the fraction of sequences with a valid fluke to form an anchor. `AnchoRNA` takes two arguments: a file containing the sequences and a configuration file (default name: `anchorna.conf`), see Fig. 2, that sets the parameters and specifies information such as the annotation of coding sequence and the choice of guiding sequence. The sequence file is parsed by the `sugar` package, which currently supports either a `.gbk` file, including annotations of viral protein positions; or a `.gff` file with `fasta` directive. The required CDS annotations may involve additional preprocessing. This enables anchor finding in amino acid space while retaining nucleotide-level information. Future versions could integrate the ORF-finding functionality already available in the `sugar` library to aid in this step.

**Figure 2:**
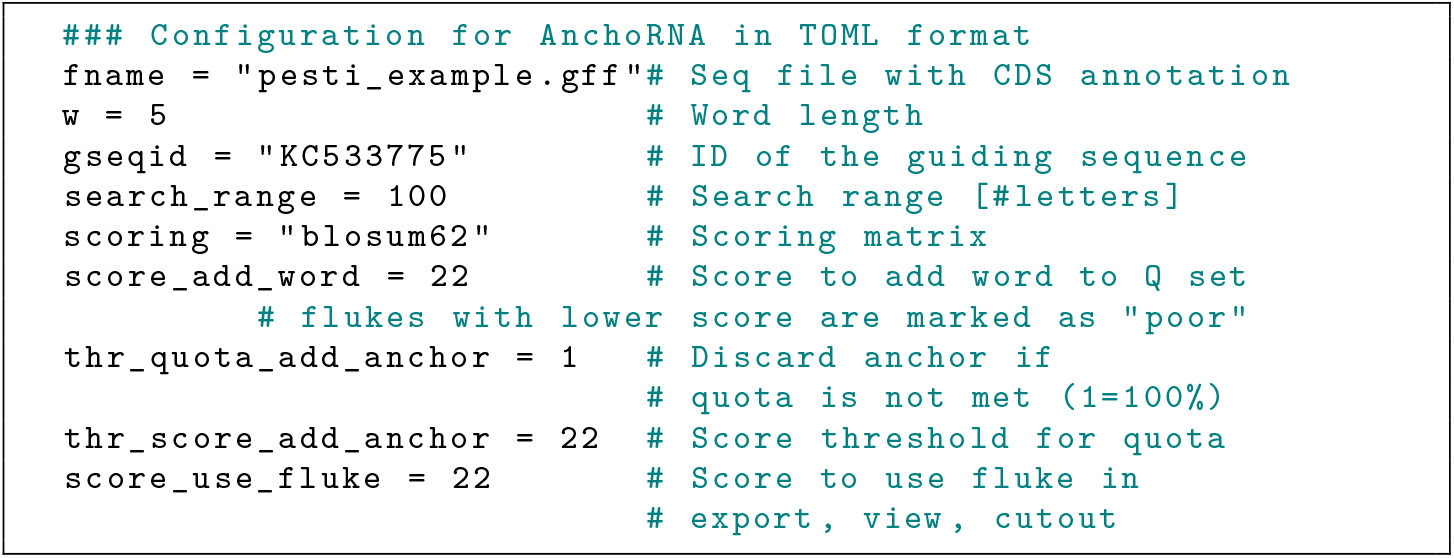
Configuration parameters for `AnchoRNA`.

The `AnchoRNA` algorithm is executed by the `anchorna go` command. The resulting anchors are written to a file with a special format based on the specification of GFF. Support for the creation of the configuration file is provided by running `anchorna create`. The software is distributed with an example data set comprising 55 *Pestivirus* genomes, allowing the user to recreate all examples shown in this contribution. In addition, a detailed tutorial is provided on the GitHub landing page (Eulenfeld, 2024).

**Default parameter settings** have been chosen to work well with this, and similar, data sets. The default length for the sliding window and initial anchors for protein sequences is *w* = 5 amino acids. The size of the query sequence can be increased for data with a higher conservation level. For Hepatitis C Viruses genomes, for example, we recommend *w* = 8. Applying `AnchoRNA` directly to nucleotide sequences, such as 5’ UTRs and 3’ UTRs, also requires larger values. We found 10 ≤ *w* ≤ 15 to work well in such a scenario. The size of `search_range` is set to *L* = 100 by default. The length of the interval in which flukes are searched for is 2*L* + *w*. Larger values are likely of use for input sequences with large length variations and for nucleic acid sequences. A potential anchor becomes a fixed anchor if a fraction of `thr_quota_add_anchor` (default: 100%) of the flukes score above *σ*_add_anchor_ (`thr_score_add_anchor`, default 22 for protein sequences scored with the `BLOSUM62` similarity matrix). Reducing the `thr_quota_add_anchor` parameter makes it possible to find anchors that are present in a subset of input sequences only. While this can be useful in the presence of larger deletions in some sequences, it makes the results strongly dependent on the choice of the guiding sequence. It is beneficial to select the guiding sequences as the longest sequences in such cases. It should be kept in mind that threshold score, window length *w*, and scoring matrix options are dependent on each other. The default values were chosen so that the product of the word length and the identity score of a single *common* letter in the `BLOSUM62` matrix is slightly smaller than the score threshold.

### General workflows

The current implementation of `AnchoRNA` supports input data comprising up to a few hundred viral genomes. For well-studied viruses, much larger sets of genomic sequence are available, many of which are then identical or near similar. It this case it is advisable to first perform a clustering by sequence similarity as described e.g. by Triebel et al. (2024, 2025). Sequences within a cluster are sufficiently similar to be aligned globally without the need of anchors. A representative or the consensus sequence of each cluster then serves as input for an `AnchoRNA`-based alignment. If necessary the alignment of the sequences within a cluster can also be performed with the help of `AnchoRNA`, which can be expected to produce a much denser set of anchors within clusters than among cluster representatives. The `AnchoRNA` software implements not only the `AnchoRNA` algorithm but also provides several additional functionalities to inspect and modify the anchor data, see Fig. 3 and Fig. S1 in the appendix. The `anchorna view` command allows the user to inspect the anchors in Jalview. Anchors can be overlaid on raw sequences or a preliminary alignment, facilitating the identification of erroneous anchors by visual inspection. The `anchorna combine` command can be used to manually delete erroneous anchors or select anchors. It can also be used to combine multiple anchor files by adjusting positions to each other. This may be necessary when using `AnchoRNA` in an iterative manner to search for anchors in “empty” regions of the sequence set where no anchors were identified in the initial, genome-wide run, see Fig. S1 in the appendix. The necessity to combine multiple anchor files can also occur, when anchors are determined separately in different coding sequences of the same viral genome. This strategy has proved useful for *Filoviridae*. `anchorna cutout` can be used to create, for each virus, a subsequence defined by locations relative to 5’ and 3’ anchors. The location strings must be from the group of strings defined by the regular expression (A?\d+|ATG|*|start|end)([<^>])?([+-]\d+)? (case is ignored). The regular expression consists of three groups: The first group gives the anchor number. Start and stop codons, as well as start and end of sequences, can also be used as anchors. The optional second group defines the location inside the anchor. It can be the 5’ or 3’ end, or the center. By default, the 5’ or 3’ end is used for anchors at the 5’ or 3’ end of the cutout region, respectively. The third group defines an optional constant character offset. For example the location A1>+6 is six characters behind the 3’ end of the second anchor. Sequences can be cut in such a way that the original sequences can be reconstituted using the `sugar merge` command provided by the `sugar` package (Eulenfeld, 2025b). This makes it possible to use `sugar merge` also to concatenate subalignment files output by alignment tools and generate the full alignment. This facility is useful in particular when alignment tools are used that do not natively support anchors and thus have to process the sequences between the anchors independently. Several alignment tools allow to input anchor files directly to guide the alignment process. The `anchorna export` command creates corresponding anchor files for the `Dialign` and `LocARNA` programs. The command can also be used to create `.gff` files with anchor positions in the nucleotide or amino acid sequences for each virus genome.

**Figure 3:**
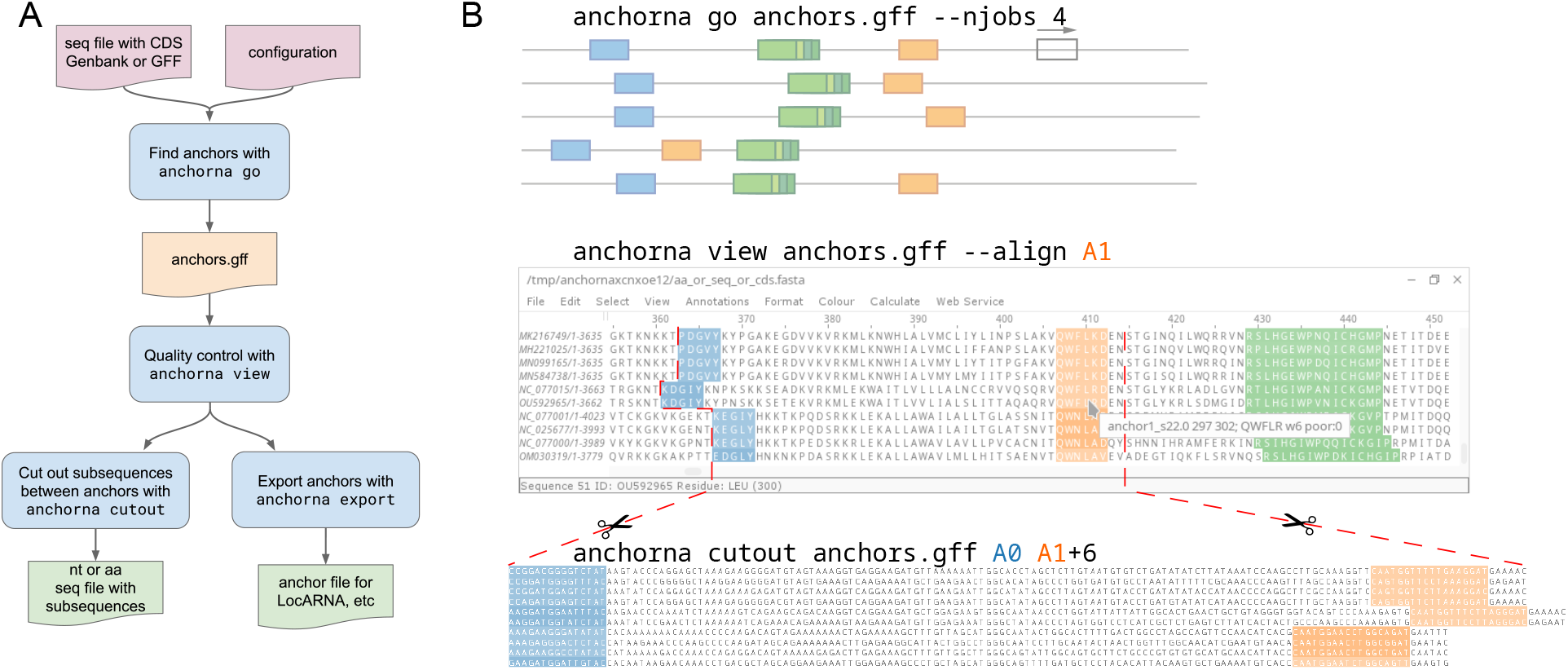
Basic `AnchoRNA` workflow. **A** The `anchorna go` command takes a sequence file and a configuration file and outputs anchors in GFF format. Anchor GFF files can be viewed with `Jalview` (Waterhouse et al., 2009) using the `anchorna view` command. The original or translated sequences can be cut at positions relative to the anchors with the command `anchorna cutout`. Anchors can be exported to different formats with the command `anchorna export`. A more advanced workflow is shown in Fig. S1 in the appendix. **B** Example usage of `AnchoRNA` commands based on the tutorial dataset. `anchorna go` is used with 4 CPUs; `anchorna view` shows anchors, here A0 (blue), A1 (orange, aligned), A2 (green); `anchorna cutout` cuts the nucleotide sequences from anchor A0 to anchor A1 and adds 6 nucleotides to the right.

## 3 Results

### 3.1 Full-genome alignment of pestiviruses

We selected the *Pestivirus* taxon as showcase application because its genomes exhibit both conserved regions that can easily be aligned using standard tools and regions that require anchors for alignment. The members of the genus *Pestivirus* (since March 2026 named *Orthopestivirus*, family *Pestiviridae*) are enveloped RNA viruses (Tautz et al., 2015). The positive-sense, single-stranded genome encodes one large polyprotein. Several members, including bovine viral diarrhea virus types 1 and 2 (BVDV-1 and BVDV-2) and classical swine fever virus (CSFV), are well-established animal pathogens (Tautz et al., 2015). Triebel et al. (2025) provided a carefully curated ground truth against which `AnchoRNA` can be compared. We used their representative dataset of 55 *Pestivirus* genomes with an approximate length of 12 kb each. These genomes were selected with the help of the `ViralClust` pipeline (Triebel et al., 2026) to ensure a well-distributed and balanced representation across the phylogenetic tree of the genus *Pestivirus*. The corresponding sequences are included as example data set with the `AnchoRNA` distribution. As a downstream application we aim to predict RNA secondary structures across the entire RNA genome alignment to evaluate the dependence of the prediction on the quality of the genome-wide alignments. We use `LocARNA` (O(*n*^2^)), a sparsified heuristic adaptation of the Sankoff algorithm (O(*n*^6^)) (Sankoff, 1985).

Using `AnchoRNA`, we translated all protein-coding regions into amino acids and identified 56 anchors across the entire genome. We identified a single erroneous anchor A8 with the `anchorna view` command, leaving 55 valid anchors. The anchors are 5–22 amino acids in length, spaced between 1 and 670 amino acids apart, see Fig. 4. For a detailed view, we selected 24 sequences (at least one from each genotype) and highlighted anchor no. 11–13, which are easy to align, and anchor 14, which only `AnchoRNA` successfully aligned (Fig. 5), see below. Triebel et al. (2025) used these anchors to split up the *Pestivirus* genome sequences into several subsequences which they fed into the `LocARNA` algorithm. We use their resulting hand-curated *Pestivirus* alignment as ground truth.

**Figure 4:**
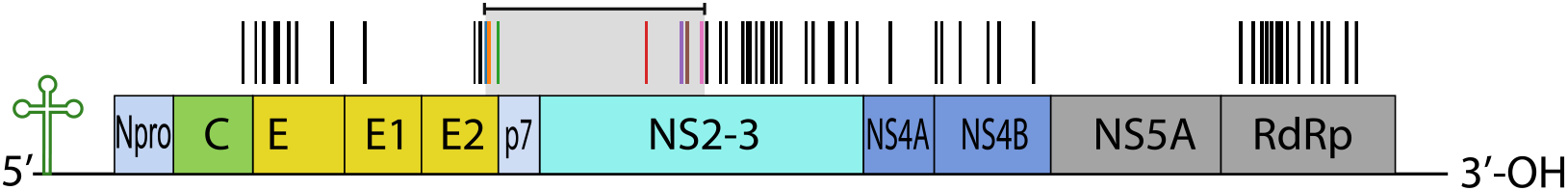
*Pestivirus* genome of around 12 kb length adapted from Viralzone (Hulo et al., 2010) with anchors indicated by black and colored vertical bars. 55 valid anchors were found using the tutorial dataset of 55 *Pestivirus* sequences. Different alignments of the genome region marked with the horizontal bar and light gray box are shown in Fig. 8. The first four colored anchors in this region are displayed in Fig. 5. The distance between neighboring anchors ranges from 1 aa to around 670 aa depending on position in the genome and sequence.

**Figure 5:**
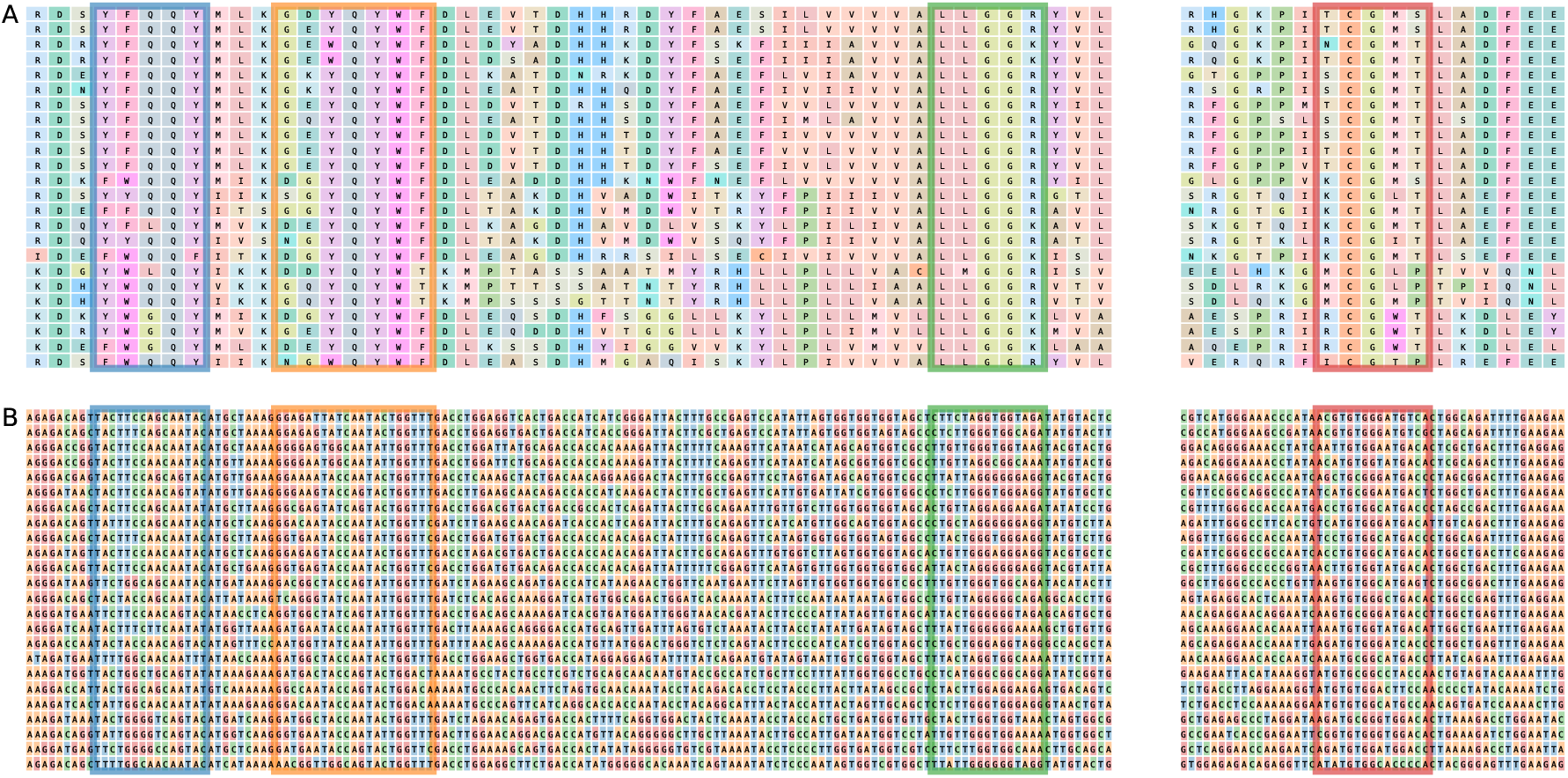
Sequences with four exemplary anchors found with `AnchoRNA` for the *Pestivirus* dataset. For visualization purposes only 24 sequences (each genotype represented) are displayed, with 1,000-1,600 nt omitted (white space). The four anchors correspond to anchor 11-14, which are displayed with the same colors in Fig. 4 and Fig. 8. `AnchoRNA` works on the amino acid level by default (**A**) and the anchors are transferred to the nucleotide level only afterwards (**B**).

### 3.2 Comparison and evaluation of different anchor-finding tools

We compare `AnchoRNA` with three tools for identifying conserved regions across multiple nucleotide sequences: `MS4`, `Murasaki`, and `MEME`. These methods were selected to represent different approaches to identifying sequence similarities and candidate anchors. We focus on methods that can derive candidate anchors from a collection of sequences, rather than tools that only report pairwise matches and require a separate procedure to integrate these matches into multi-sequence anchors. Consequently, pairwise anchor-finding tools such as `CHAOS` and `MUMmer` are not included, since developing and evaluating an additional aggregation algorithm is beyond the scope of this study. `MS4` was included because it was specifically developed to identify multiple local similarities (Corel et al., 2010; Pitschi et al., 2010). To ensure that the identified anchors were supported across the complete sequence set, we configured MS4 to consider only partitions containing all input sequences. We set the window parameter *N* to 2–60, following the settings used by the developers in their analysis of hepatitis C virus genomes. `Murasaki` was included because it was explicitly designed to identify anchors across multiple sequences simultaneously (Popendorf et al., 2010). It uses spaced-seed patterns to identify matching sequence regions, extends matching seeds, and combines compatible overlapping anchors. We selected the pattern `110110110110110` which matches 5 codons while allowing a mismatch at the third position of each codon, thereby accommodating synonymous substitutions at wobble positions. We experimented with different patterns, this pattern gave the most valid anchors. `MEME` was included as a motif-discovery-based approach (Bailey and Elkan, 1995; Bailey et al., 2006). It identifies statistically significant sequence motifs in collections of nucleotide sequences. We selected the occurrence model that specifies exactly one occurrence of the motif in each input sequence. We allowed motif widths of up to 50 nucleotides. The resulting motifs capture conserved nucleotide patterns while accommodating variation at other positions, which may include codon wobble positions. We also evaluated one variation of `MEME`, hereafter referred to as `MEME_aa`, that operates on translated amino acid sequences. We translated all coding sequences and applied `MEME` to identify motifs with width ranging from 5 to 20 amino acids. We then mapped the resulting motif positions back to the nucleotide sequences.

For the evaluation of the different anchor methods we reduced the *Pestivirus* dataset from 55 to 50 sequences. On this reduced dataset of 50 sequences, `AnchoRNA` identified the same 55 valid anchors as in the full dataset of 55 sequences. We evaluated the anchors identified by each method using the hand-curated *Pestivirus* multiple sequence alignment as a reference. Compatible overlapping anchors were merged in a first step. An anchor was considered valid if either start or stop positions of all flukes in one anchor aligned in the reference alignment and the anchor contained no more than 20 gaps over all sequences. For the evaluation we used a single core of an AMD EPYC 9754 128-Core Zen 2.25 GHz machine. Table 1 reports runtime, the number of merged anchors, the proportion of valid anchors, the resulting precision, and summary statistics (minimum, median, maximum) for anchor lengths and distances between neighboring anchors. The runtimes of `Murasaki` and `AnchoRNA` ranged from seconds to minutes, whereas `MEME`, `MEME_aa`, and `MS4` required several hours to complete. `MS4` identified only three anchors, all of which were valid. `Murasaki` and `MEME` achieved similar precision (approximately 13 %), but `MEME` identified more valid anchors (13 compared with 7 for `Murasaki`). `AnchoRNA` and `MEME_aa` identified a much higher number of valid anchors than the other methods (55 and 60, respectively); however, `AnchoRNA` achieved substantially higher precision (98%) than `MEME_aa` (65%). For each valid `AnchoRNA` anchor, we assessed whether it overlapped with anchors identified by the other tools. As shown in Figure 6, all valid `AnchoRNA` anchors overlapped with anchors identified by the other tools, except for `MEME` and `MEME_aa`. Furthermore, `MEME_aa` identified 21 anchors that were not detected by `AnchoRNA`, whereas `AnchoRNA` identified 9 anchors that were not detected by `MEME_aa`. 2 anchors identified by `MEME_aa` were also found by `MEME`. Figure 7 illustrates the distribution of anchors identified by the different tools across the *Pestivirus* genome. `MEME_aa` and `AnchoRNA` produce the densest anchor distributions, with maximum distances between neighboring anchors of 1 855 nt and 2 094 nt, respectively. Overall, `AnchoRNA` outperforms the other methods in terms of precision and the number of valid anchors identified. Although `MEME_aa` detects more valid anchors, the large number of unaligned anchors reduces its practical applicability. Since `MEME` was not specifically designed for anchor detection, applying a dedicated post-processing algorithm to filter the identified motifs could substantially improve the precision of `MEME_aa`.

**Table 1:** Comparison of anchor-finding methods on the *Pestivirus* dataset, including basic statistics on anchor lengths and distances between neighboring anchors.

|  | runtime (s) | anchors | valid | precision | length (nt) |  |  | distance (nt) |  |  |
|---|---|---|---|---|---|---|---|---|---|---|
|  |  |  |  |  | min | med | max | min | med | max |
| AnchoRNA | 160 | 56 | 55 | 98 % | 15 | 21 | 66 | 3 | 790.5 | 2 094 |
| MS4 | 22 000 | 3 | 3 | 100 % | 31 | 34 | 43 | 98 | 1 990 | 10 607 |
| Murasaki | 2.8 | 57 | 7 | 12 % | 15 | 60 | 456 | 34 | 1 539 | 5 439 |
| MEME | 24 000 | 100 | 13 | 13 % | 41 | 50 | 50 | 4 | 377.5 | 4 888 |
| MEME_aa | 4 600 | 92 | 60 | 65 % | 36 | 60 | 144 | 3 | 33 | 1 855 |

**Table 2:**
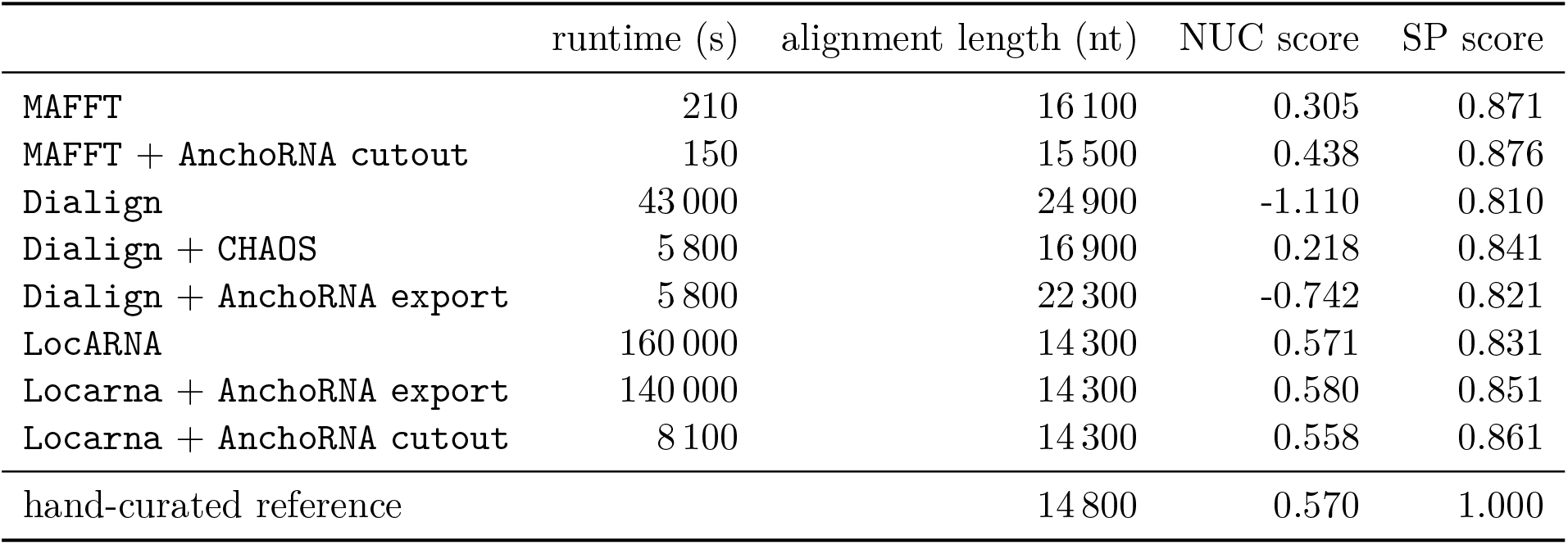
Comparison of alignment methods with and without guidance by anchors of the curated *Pestivirus* sequences. NUC scores are mean scores per alignment column calculated using the `NUC4.4` matrix, with a maximum possible score of 5. SP scores represent the fraction of aligned residue pairs that agree with the reference alignment. Runtimes for AnchoRNA-guided alignments exclude anchor-finding time.

**Figure 6:**
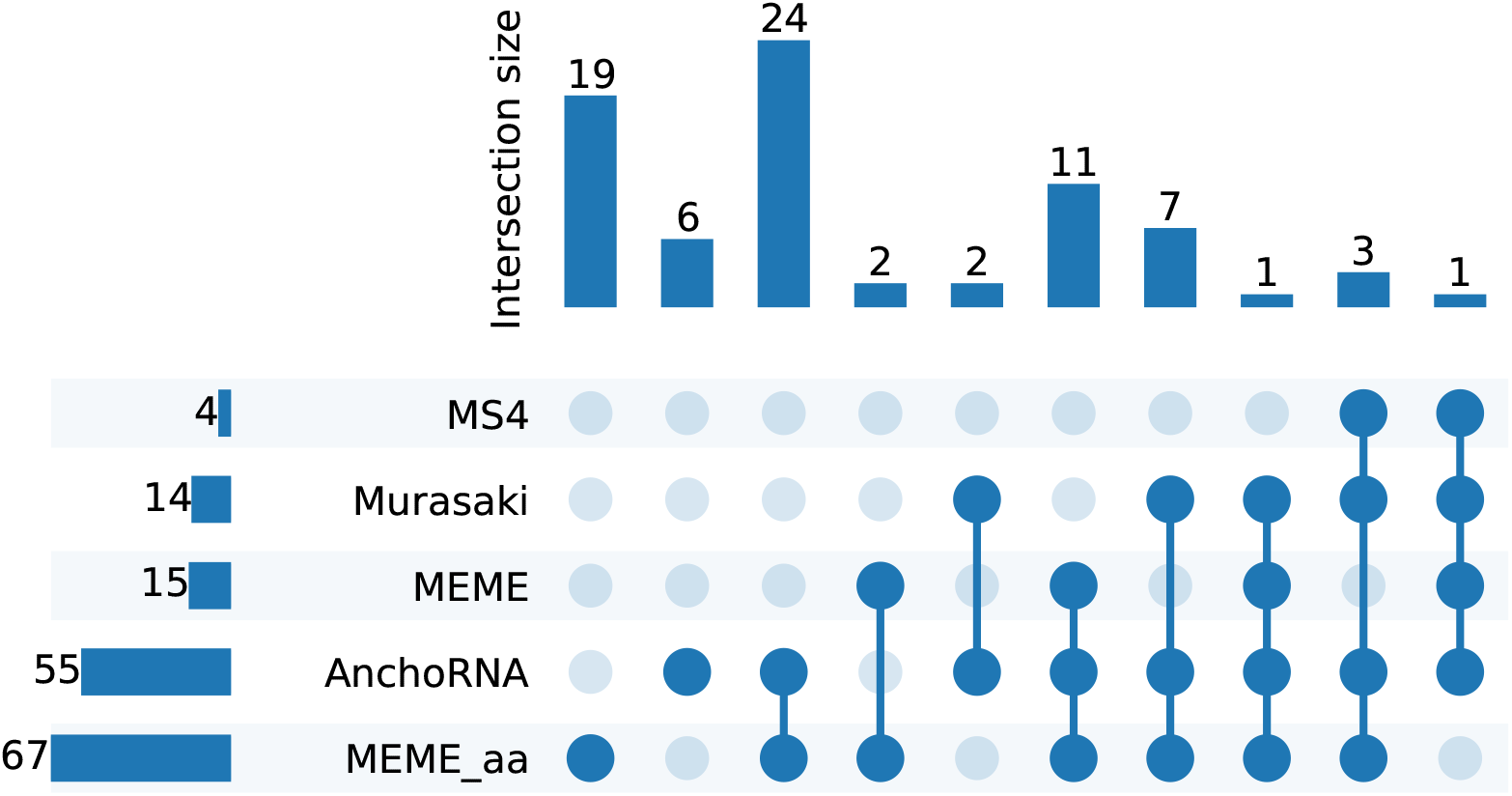
Upset plot of the number of `AnchoRNA` anchors overlapping with anchors found by other tools, the number of anchors found only by both `MEME` and `MEME_aa` and of the number of anchors solely found by `AnchoRNA` or `MEME_aa`.

**Figure 7:**
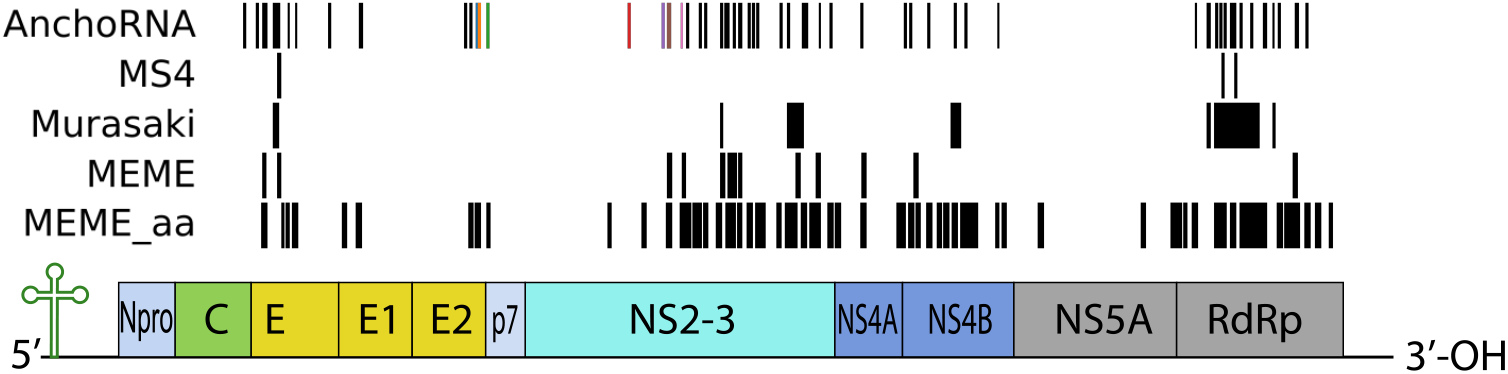
*Pestivirus* genome of around 12 kb length adapted from Viralzone (Hulo et al., 2010) with valid anchors found by different tools indicated by black and colored vertical bars.

### 3.3 Impact of anchors on different alignment tools

To validate our anchors independent from the validation in the previous section, we first overlaid them on a `MAFFT` alignment using the `anchorna view` command. All anchor regions were aligned by `MAFFT`, except for the erroneous anchor mentioned above and anchor A14. The region between anchors A13 and A15 appears difficult to align due to mutations and large deletions and insertions. This region spans between 430–700 amino acids, and we will focus on it below.

We compared the impact of `AnchoRNA` anchors on several alignment tools. We chose `MAFFT` (FFT-NS-i algorithm) for its popularity and widespread use, and `MAFFT` with `AnchoRNA` to assess the effect of anchor-based refinement. `Dialign` was included because it is well-suited for handling insertions and deletions, while `Dialign` with `CHAOS` has been described to utilizes pairwise anchors to enhance alignment quality. We evaluated `Dialign` with `AnchoRNA` to integrate our anchor-based approach into the `Dialign` workflow. Finally, we also employed `LocARNA` in its standard form, alongside `LocARNA` guided by `AnchoRNA` anchors, and `LocARNA` applied to subsequences. The choice of using the computationally expensive tool `LocARNA` reflects the growing importance of analyzing RNA secondary structures across entire viral genomes. Our method assumes that homologous regions can be aligned in a consistent linear order across the input sequences. Consequently, it is best suited to datasets in which genomic rearrangements, duplications, or other structural variations do not disrupt this collinearity. It is not intended to replace whole-genome alignment methods specifically designed to represent structural differences between genomes.

The various tools necessitate different adaptations for alignment. We used the `Dialign` web service^1^, which cannot process more than 50 sequences. Therefore, the dataset was reduced accordingly to 50 sequences (see above). For `AnchoRNA`+`Dialign`, we used the `anchorna export` command to export the anchors A14 and A41, which were not aligned by vanilla `Dialign`, along with the surrounding anchors, into the `Dialign` anchor file, which serves as a constraint for the `Dialign` alignment process. Note that not all 55 valid anchors were exported because Dialign could not handle so many constraints, which resulted in unaligned anchor regions. For `AnchoRNA`+`MAFFT`, we only chose anchor A14 as constraint because Vanilla `MAFFT` aligned all other valid anchor regions out of the box. We utilized the `anchorna cutout` command to split the genome sequences into two subsets at anchor A14, as `MAFFT` does not provide an option to directly incorporate anchors. This allowed us to force `MAFFT` to align the anchor, after which we concatenated the resulting alignments with the `sugar merge` command. For `AnchoRNA`+`LocARNA`, we exported all valid anchors, to the `LocARNA` anchor format using the `anchorna export` command. These anchors were then utilized as constraints for `LocARNA`, enabling more accurate alignment by guiding the alignment process based on the identified anchor points. In another approach, we split the genome at the anchors and fed each subsequence set into `LocARNA`, subsequently concatenating the resulting alignments. Detailed commands to reproduce the analyses are available in our repository : https://github.com/rnajena/anchorna_paper (Eulenfeld, 2025a).

For our tests, we use a standard PC with four 3.6 GHz i7-7700 cores, each with two threads, and 67 GB of memory. Two threads were used with `LocARNA`, while `MAFFT` used only a single thread. Fig. 8 displays parts of the resulting alignments for the various alignment approaches for *Pestivirus* genomes between `AnchoRNA` anchor no. 11 and 17. The most compact alignment-part, produced by `LocARNA`, measures ≈2,700 nt in length, while the widest alignment, optimized for longer insertions using `vanilla Dialign`, extends to ≈6,980 nt. Notably, while this example highlights a single anchor region at the nucleotide level that is not aligned by standard multiple sequence alignment methods, in more complex real-world scenarios, this issue typically affects numerous such regions. To measure a precise alignment score, we used Todd Lowe’s `NUC4.4` scoring system^2^, which assigns +5 for identical letters, -4 for different letters, additionally we set -15 for a new gap, and -1 for gap extensions. The choice of scoring system is not unique, and the resulting score should not be interpreted as a direct measure of biological alignment accuracy. Moreover, finding an optimal multiple sequence alignment under such a scoring scheme is NP-hard (Wang and Jiang, 1994). We calculate the alignment score for each sequence pair, compute the mean of these scores, and divide by the constant median length of the 50 sequences (12 296.5). This results in a mean alignment score per alignment column, with a perfect score of 5. We complement the mean alignment score with a reference-based sum-of-pairs (SP) score, which measures the fraction of residue pairs aligned together in both the evaluated alignment and the hand-curated *Pestivirus* reference alignment. The SP score ranges from 0 to 1, with 1 indicating agreement with the reference alignment. Tab. 2 compares various alignment methods for the 50 *Pestivirus* genomes, both with and without anchor guidance. Without anchors, `MAFFT` produced a reasonable alignment with a length of 16.1 kb and a mean score of 0.30. In contrast, `Dialign` generated a much longer alignment being gap-rich of 24.9 kb length, resulting in a low mean score of -1.11. `LocARNA` produced the alignment with the highest score of 0.57 and an alignment length of ≈14.3 kb. When anchor guidance by `CHAOS` was introduced, `Dialign` benefits substantially, achieving a score of 0.22 with a more manageable alignment length of 16.9 kb. This result can be attributed to the fact that `CHAOS` calculates anchors specifically for pairs of sequences. In general, these pairwise anchors do not necessarily determine gap-less local alignment blocks and thus, do not induce a partition of the global multiple alignment problem. Pairwise anchors therefore cannot be coupled with arbitrary alignment algorithms. Instead, the alignment algorithm itself must be capable of handling pairwise alignment constraints. No such feature is available for `MAFFT` and `LocARNA`. We note that pairwise anchors need to satisfy non-trivial consistency conditions, which have been specifically designed into `Dialign` (Morgenstern et al., 2006).

**Figure 8:**
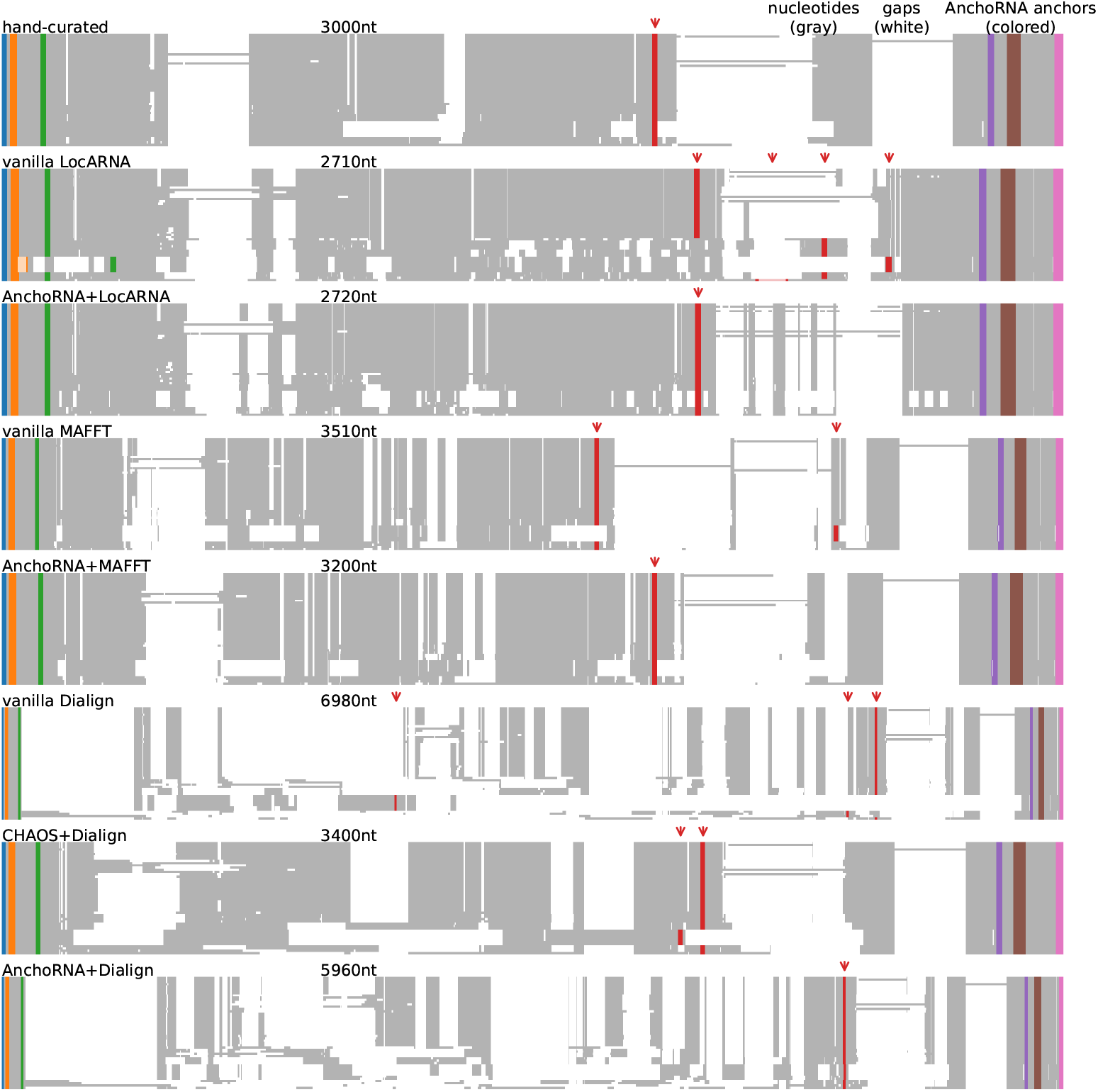
Part of the alignments produced by different tools with and without guidance by anchors. Anchors calculated by `AnchoRNA` are displayed over each alignment with colored bars. The positions of anchor 14 are marked with red arrows. Length of the displayed parts of alignments rounded to 10 nt are indicated. Each panel corresponds to one cell in Tab. 2. The guidance by anchors decreases alignment length for most tools.

All alignment methods benefited from the guidance of `AnchoRNA` anchors. `MAFFT` could be improved to a mean score of 0.44 and an alignment length of 15.5 kb. `Dialign` in combination with `AnchoRNA` led to an improvement with a score of -0.74 and an alignment length of 22.3 kb, but this was not as efficient as the results obtained with `CHAOS` anchors, which use a larger number of pairwise anchors. Note that we could have further improved the `AnchoRNA` +`Dialign` alignment by searching and incorporating anchors that span a subset of sequences in the crucial region between anchors A13 and A15. Finally, we input `AnchoRNA` anchors into `LocARNA`, which resulted in a slightly improved alignment score of 0.58, however, it still had an extensive runtime of around two days. Therefore, we also ran `LocARNA` on subsequences based on the calculated anchors, which yielded a slightly lower alignment score of 0.56 and an alignment length of 14.3 kb, but with a more reasonable runtime of 2 h. This result has a similar score as the manually curated ground truth alignment by Triebel et al. (2025), which achieved an alignment score of 0.57 and an alignment length of 14.8 kb. The manually curated alignments are partially based on pure secondary structure conservation, which leads to less optimal sequence alignments. Consequently, while the alignment score is similar, it is considered to more accurately reflect the evolution of the virus family. SP scores ranged from 0.810 for `DIALIGN` to 0.876 for `MAFFT` guided by `AnchoRNA`, indicating varying levels of agreement with the hand-curated reference alignment. Overall, `MAFFT`-based alignments showed the highest agreement, followed by anchor-guided `LocARNA`, anchor-guided `DIALIGN`, vanilla `LocARNA`, and vanilla `DIALIGN`. Although the differences in SP scores were relatively small, the lower NUC scores for `DIALIGN` suggest that it produces alignments with more gaps than the other methods (Figure 8).

## 4 Conclusion

The alignment of complete viral genomes is challenging due to the complexities associated with long sequences, extensive insertions and deletions, and high sequence divergence. Our study demonstrates that `AnchoRNA` effectively addresses these challenges by identifying conserved regions, or anchors, within the coding sequences, demonstrated at the example of pestiviruses. Utilizing a dataset of 55 representative genomes, `AnchoRNA` identified 55 valid anchors that are used for guiding the alignment process. The incorporation of these anchors led to improvements across tested alignment tools, highlighting the effectiveness of `AnchoRNA` in enhancing alignment quality, especially in viral genomes. Even in cases where alignments are not improved in terms of score, forcing the sequences to align in conserved anchor regions leads to a biologically more meaningful alignment (Triebel et al., 2025).

A limitation of this study is that AnchoRNA was evaluated on genomes from a single genus, and its anchors also assisted manual curation of the reference alignment. This may introduce bias into the evaluation. Future work will focus on applying AnchoRNA to additional viral genera and generating high-quality genome alignments for a broader range of viruses.

While not the primary focus of our tool, `AnchoRNA` is capable of identifying anchors in both amino acid and non-coding sequences without requiring translation. Though, effectiveness of the tool will drop when sequence conservation is low and structural anchors are desired. Its rapid processing time makes `AnchoRNA` a viable option for researchers requiring timely results without sacrificing the accuracy of alignment. The ability to handle viral genomes and other highly variable sequences sets `AnchoRNA` apart from existing methods, making it a valuable addition to the MSA toolkit.

Furthermore, the abundance and distribution of anchors identified by `AnchoRNA` represent a significant contribution for primer design across entire virus families. The conserved nature of these anchors ensures effective amplification of genetic material, which is crucial for diagnostics, epidemiology, and virology research. Therefore, integrating `AnchoRNA` into alignment and primer design processes not only enhances alignment quality but also makes a meaningful contribution to the broader field of viral research and management.

## Availability and requirements

**Project name** AnchoRNA

**Project homepage** : https://github.com/rnajena/anchorna

**Releases** are distributed via PyPI and archived at Zenodo (Eulenfeld, 2024)

**Operating systems** Linux, MacOS, Windows

**Programming Language** Python

**Other requirements** Python 3.11+, rnajena-sugar

License MIT

**Restrictions to use by non-academics** MIT license

## Declarations

### Ethics approval and consent to participate

Not applicable.

### Consent for publication

Not applicable.

### Availability of data and materials

The necessary code, data, and results to reproduce this study are available on GitHub at : https://github.com/rnajena/anchorna_paper and archived on Zenodo (Eulenfeld, 2025a).

### Author contributions

M.M. and T.E. conceptualized the tool. T.E. developed and evaluated `AnchoRNA`. T.E. and S.T. tested the software. T.E. and M.M. wrote the manuscript. P.F.S and T.E. revised the manuscript.

## Acknowledgments

We thank Daria Kubrak for drawing the scheme of the method.

## Appendix Pseudocode of the AnchoRNA algorithm

### Algorithm 1

Anchor-finding algorithm underlying the `anchorna go` command

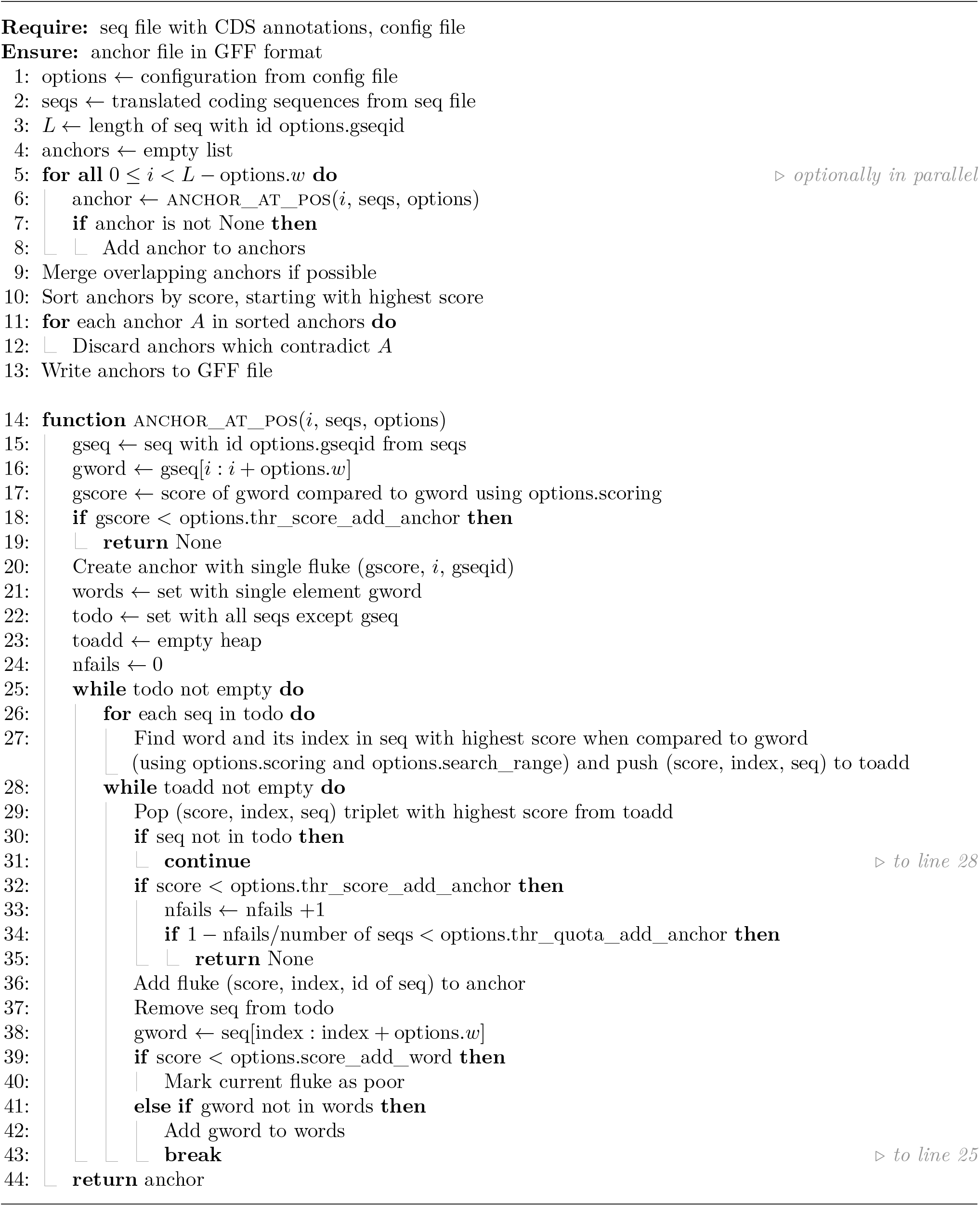

## Appendix Extended workflow

Fig. S1 shows an extension of the workflow chart in Fig. 3. It includes commands to refine the anchor search in specific regions of the virus genome.

**Figure S1:**
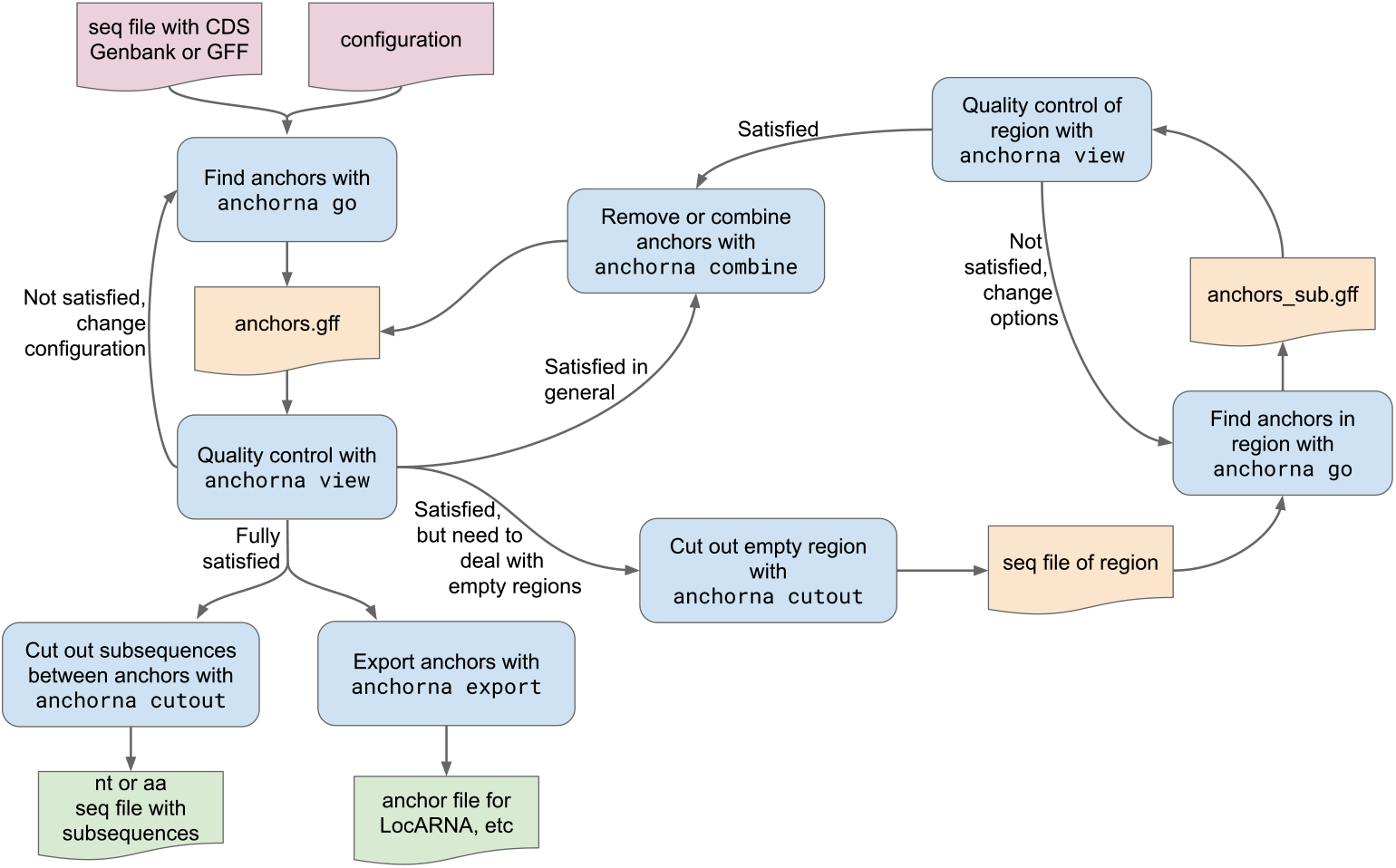
The place of the different `AnchoRNA` commands in the general workflow. The output of several commands, represented by orange data boxes, may be directly passed to other commands using a pipe.

## Footnotes

1 https://dialign.gobics.de/, last accessed 07 Apr 2025

2 ftp://ftp.ncbi.nih.gov/blast/matrices/, last accessed on 21 Dec 2024, also distributed with the `sugar` package

